# Particulate Contamination of Human Placenta: Plastic and non-plastic

**DOI:** 10.1101/2024.03.01.583005

**Authors:** Rewa Zurub, Shannon Bainbridge, Luna Rahman, Sabina Halappanavar, Michael G. Wade

## Abstract

Recent evidence indicates that the human womb is contaminated with a variety of particulate contaminants. Microplastics (MPs, tiny plastic particles (≤5 mm) generated by the breakdown of larger plastic products in the environment) accumulation in human placenta has recently been described. In addition, recent evidence has correlated the number of air pollution particulates in term placentas to the loading of these particles in dust from the gestational parent home. The current study sought to characterize the accumulation of plastic and non-plastic particles (NPP) within the term human placenta. Placenta tissues were collected from healthy, singleton pregnancies following vaginal (n=5) and caesarean section (n=5) deliveries at a tertiary care centre located in an urban Canadian city (Ottawa, ON), with particles detected and characterized by Raman micro-spectroscopy. Both plastic and non-plastic particles were identified in all placentas examined, with an average of 1 ± 1.2 MPs /g and 4 ± 2.9 NPP /g of tissue. Similar tissue concentrations of MPs and NNP were identified in all regions of the placenta (basal plate, chorionic villous, chorionic plate), and did not differ according to mode of delivery. MPs ranged in size (2 - 60 μm), with the most abundant MPs being polyethylene (PE), polypropylene (PP), polystyrene (PS) and polyvinyl chloride (PVC). The most abundantly identified NPP were carbon, graphite, and lead oxide. Collectively, these results demonstrate the accumulation of foreign particles, including MPs, throughout the human placenta. Given the vital functions of the placenta in supporting fetal growth and development, and a potential for MPs to induce toxicity, further investigations into the potential harmful effects of these environmental toxicants on maternal and fetal health is warranted.

**Graphical Abstract:** 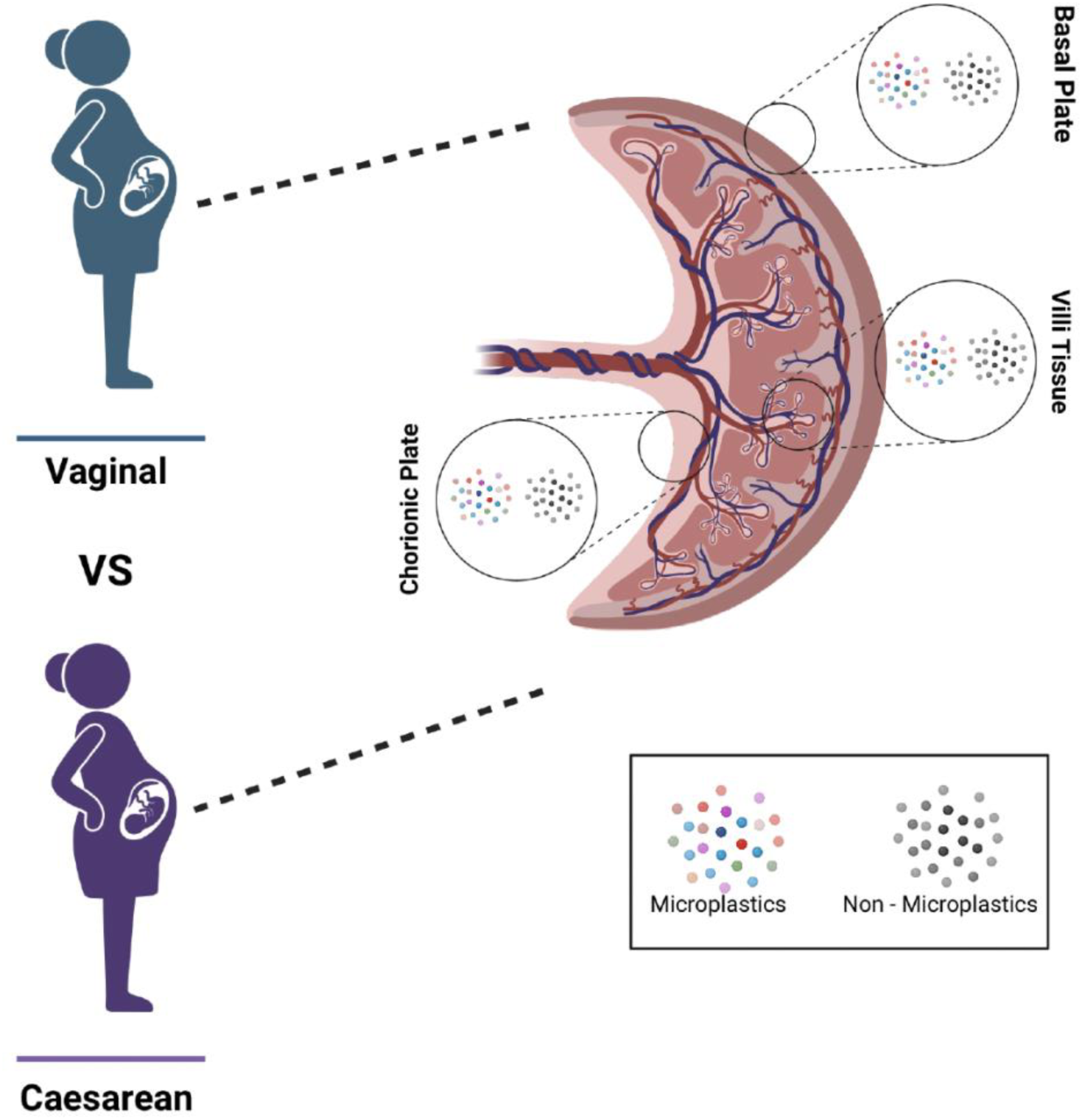

**Highlights:**

- Both plastic and non-plastic foreign particles were observed in term placentas from Canadian women
- Placenta contamination with most of the non-plastic particle types observed has not previously been reported.
- Particles were observed in similar frequency in placenta regions corresponding to maternal or fetal circulation suggesting that particles pass unhindered through the placenta.

## 1. Introduction

In the past few years, the environmental presence of microplastics (MPs) has come to the fore as a major concern for human exposure and potential health consequences. Intense plastic manufacturing and use, coupled to poor waste management, has led to an environment overwhelmed with billions of tons of plastic (Geyer et al., 2017; Jambeck et al., 2015). Plastic material abandoned in the environment leads to fragmentation and degradation (Andrady, 2017). This process yields smaller plastic fragments referred to as MPs. MPs range in size, from 1µm to 5mm (Hartmann et al., 2019), and are found ubiquitously in the environment, including waterways (Eriksen et al., 2013; Gaylarde et al., 2021) soil (Ramos et al., 2015) tap water (Kosuth et al., 2018), household dust (Zhang et al., 2020) and food products (Afrin et al., 2022; Diaz-Basantes et al., 2020; Kosuth et al., 2018).The ubiquity of these contaminants across these media argue that widespread human exposure is inevitable.

Numerous investigations, encompassing both human and animal studies, have established that particulate – primarily MPs - have the capacity to enter the human body and collect within tissues and organ systems (Amato-Lourenço et al., 2021; Baeza-Martínez et al., 2022; Horvatits et al., 2022; Ibrahim et al., 2021; Jenner et al., 2022; Ozawa et al., 1986; Pauly et al., 1998) . MPs have been detected in various human biological matrices, including, human blood (Leslie et al., 2022), urine (Massardo et al., 2024; Pironti et al., 2022), sputum (Huang et al., 2022), breast milk (Ragusa et al., 2022b), semen (Montano et al., 2023)), diverse body fluids (collected from cysts, synovia or organ effusions, etc; (Guan et al., 2023) and fecal matter (Schwabl et al., 2019). Furthermore, these investigations have illuminated the distribution of MPs within various human organ systems encompassing the lungs (Amato-Lourenço et al., 2021; Baeza-Martínez et al., 2022; Jenner et al., 2022; Ozawa et al., 1986; Pauly et al., 1998), spleen (Kutralam-Muniasamy et al., 2023), liver (Horvatits et al., 2022), kidney (Kutralam-Muniasamy et al., 2023), and colon (Ibrahim et al., 2021). These discoveries are causing concern regarding the potential ramifications of environmental contaminants, specifically MPs, on the health and development of the human fetus (Jummaat et al., 2021) .

In addition to MPs, humans are ubiquitously exposed to a diversity of other NPPs, both anthropogenic and natural, from many other sources (Buzea et al., 2007; Griffin et al., 2017). Recent studies have found that human tissues contain diverse NPPs. For example, ambient black carbon and other foreign particles have been discovered in human blood, as well as other tissues and organs including the heart(Calderón-Garcidueñas et al., 2019b), brain (Calderón-Garcidueñas et al., 2019a; Maher et al., 2020), ovaries (Bongaerts et al., 2023), colon (Gatti, 2004) lungs (Campion et al., 2018), liver and kidney (Gatti and Rivasi, 2002; Locci et al., 2019). However, no study to date has attempted to compare the relative levels of these diverse particle types in any tissue.

The placenta is a transitional but vital organ of pregnancy, responsible for all maternal-fetal exchange required to support fetal growth and development. Several recent studies identified the presence of MPs within the human placenta (Amereh et al., 2022; Braun et al., 2021; Liu et al., 2022a, 2022b; Ragusa et al., 2022a, 2021; Zhu et al., 2023) and fetal meconium ^24–26^ suggesting that MP can cross the placenta. In rodent models of gestational MP exposure, placental uptake has been described, with observed consequences for placental function and compromised fetal growth trajectories (Amereh et al., 2022). Furthermore, studies have revealed the presence of black carbon particulate matter – derived from particulate air pollution - in the human placenta (Bové et al., 2019); cord blood (Bongaerts et al., 2022), and fetal organs (Bongaerts et al., 2022). These findings provide evidence that adverse effects of particulate pollutants – especially air pollution - may exert adverse health effects via direct exposure to internal organs and even the developing human fetus (Bové et al., 2019).

While placenta may be exposed to MPs from the environment, given the heavy reliance on plastic polymer products within obstetrical health care delivery (Jummaat et al., 2021), there are additional questions surrounding the contributions of post-delivery contamination within the small observational studies reporting these findings, particularly across various delivery environments (i.e. surgical suite for C-section deliveries) (Braun et al., 2021).

The present study aimed to evaluate the accumulation of particulates in human placentas collected from healthy, pregnant Canadian women. The Raman micro-spectroscopy was used to specifically detect and characterise particle chemical composition. Detailed investigation of particulate localization across the distinct compartments of the placenta was undertaken, as well as a comparison of particulates in placentas delivered by vaginal or caesarean section (C-section), to address concerns related to delivery environment and potential post-delivery contamination. This approach allows identification of MPs and many NPPs to be characterized, thus providing a more comprehensive assessment of particulate exposures.

## 2. Methods

### 2.1. Patient Recruitment

This study received approval from Health Canada-Public Health Agency of Canada Research Ethics Board (REB 2021-033H) and the University of Ottawa Research Ethics Board (#H-03-22-7960). All recruited women had singleton, uncomplicated term pregnancies at The Ottawa Hospital General Campus Birthing Unit. Exclusion criteria for study participation included any obstetrical complication during pregnancy or delivery, C-section delivery for any reason other than breach presentation or repeat C-section delivery, multiple pregnancy, or participants who did not understand either English or French. A total of 10 patients were recruited: 5 undergoing a standard vaginal delivery and 5 undergoing an elective C-section delivery.

### 2.2. Characterization of potential sample contamination during processing

A plastic-reduced protocol was developed for both vaginal and C-section deliveries. All plastic-containing materials used during a standard vaginal or C-section delivery at The Ottawa Hospital General Campus were identified, and where available a non-plastic alterative material was identified for use during delivery. For those plastic-containing materials that could not be substituted due to institutional standard of care operational procedure (i.e., plastic surgical drape, gloves), items were sampled to determine potential post-delivery contamination signal. Briefly, in a sterile environment, the plastic-containing materials were rinsed three times with 30 ml de-ionized filtered water, with the rinsate collected into sterile glass bottles. The samples were filtered through a silicon membrane (1 μm retention limit, Microplastic Sample Preparation Kit, Thermo Fisher Nepean, ON) using glass vacuum filtration funnel. Silicon filter membranes were used to collect any MPs in these wash samples as their low optical interference and fine retention diameter (1 μm) ensured a high likelihood of identifying MPs compared with glass fibre membranes required for the more difficult to filter tissue samples (see below). Raman micro-spectroscopy was used to identify the presence and characterise any background plastic contamination, as described below (Section 2.4).

Similarly, plastics reduced conditions were developed for all tissue dissection and sample processing. All plastic materials (with the exception of nitrile gloves) were kept out of the biocontainment hood in which samples were handled and sample filtering was completed. Individuals working in this hood wore only clothing (including lab coats) made from natural fibres (cotton, wool or linen). Prior to use, all containers, sample vials, and glassware used in transporting, collecting, processing, or storing placenta or blank samples were vigorously rinsed with 50% of container volume with deionized water that was filtered through glass fibre membranes (Grade F, 0.7 μm retention limit, Cat# F4700, Sterlitech, Aubern, WA).

### 2.3. Placenta collection

After delivery, all placentas were collected under sterile conditions, placed into a sealed metal container, and transported on ice to the laboratory for processing within 1 hour of delivery. Placentas were weighed and placed into a sterile glass Pyrex tray within a biosafety hood. A metal bowl with 10ml deionized filtered water was placed in the hood to assess the presence of contaminating MPs within the hood air (Hood Blank - HB). Micro-dissections of placental tissue were carried out midway between the placental margin and umbilical cord insertion site. A total of 3 g of placenta tissue was collected from each placenta (1 g per micro-dissection site), specifically, sampling from the basal plate (maternal surface), chorionic villous tissue (maternal-fetal exchange region), and chorionic plate (fetal surface). All samples and HBs were placed in labelled low-particle glass vials and sealed. The water rinsate (100ml) used in the dissection process was collected into a 125 ml glass Wheaton bottle with phenol cap with polytetrafluoroethylene (PFTE) liner. Prior to sample collection, all vials and containers were rinsed 2 times with deionized water (50% total container volume) which was pre-filtered through fibre filter membranes (Grade F, 0.7 µm pore size, Sterlitech). All collected samples were stored at -20°C until digestion.

### 2.3 Digestion of placenta samples

The digestion of placenta samples used a previously published method (Jenner et al., 2022), with minor modifications, as described. All glassware used to process tissue for MP isolation was washed, rinsed 2X with deionized water then rinsed with filtered deionized water and sealed with caps (vials) or rinsed tinfoil (flasks) prior to use. Vials containing placenta tissue were thawed at room temperature, rinsed with filtered water, and placed within a biocontainment hood. Cleaned and rinsed foil-sealed glass Erlenmeyer flasks (500 ml) were weighed to the nearest 10 mg and placed into the biocontainment hood. For each micro-dissected sample from each placenta, 1 g of placenta tissue was placed into a flask, resealed, and reweighed. The flask was then returned to the hood and 100 ml of filtered (Grade F) 30% H_2_O_2_ (>95%, Thermo Scientific, Nepean) was added. The resealed flasks were then placed in a shaking incubator at 55°C for 11 days at 100 rpm. After the first 5 days of incubation an additional 50ml of filtered (Grade F) 30% H_2_O_2_ was added to the digest sample, which was then placed back in the incubator for an additional 6 days. For each batch of samples digested, a Procedural Blank (PB) was also processed. The PB flask was treated identically to all other flasks except that no tissue was added.

At the end of the incubation period, digests and PBs were filtered through glass fiber membrane (Whatman Grade GF/D, 2.7 µm pore size, Cytivia, Thermo Fisher, Nepean), using a glass vacuum filtration apparatus. Glass filtration assemblies and clamp were pre-warmed to 55 °C. Digests were poured through the membrane, followed by 10 ml of filtered (Grade F) deionized water warmed to 55°C to remove any remaining H2O2 residue. The glass fiber membranes were then individually placed in separate, clean glass petri dishes with unique identification and left in the plastic-free biosafety hood to dry.

### 2.4. Characterization of MPs by Raman Micro-spectroscopy

The Raman micro-spectroscopy method is previously published (Rahman et al., 2021). In brief, dried glass fiber membranes were examined using an integrated imaging system that couples an Enhanced Dark Field (EDF) optical microscope (CytoViva, Inc. Auburn, AL, USA) with a confocal Raman imaging system (Horiba Scientific XploRa Plus, Japan). The LabSpec 6 software suite (Horiba Ltd, Piscataway, NJ, USA) was used to acquire the Raman spectral information and KnowItAll spectral database (Bio-Rad Laboratories Inc, Hercules, CA, USA) was used to chemically identify e MPs present and other particulate matter. Membranes were visually scanned under a 10 X objective in a grid fashion to locate all particles adherent to the glass fiber membranes. Upon finding a particle, a brightfield image of the particle was captured (under 50X power). Using the 10x or 50x objective, the centre of the particle was excited at 532 nm wavelength with 1-10 mW laser power and 300nm slit and the Raman spectrum collected with 5 – 25 accumulations and 10-20s exposure duration. The collected spectrum was then queried using KnowItAll software, which compares the spectrum to a database of Raman spectra (incorporating the SLOPP and SLOPP-E microplastics databases) (Munno et al., 2020) and returns the best spectral match to predict the particle composition.

### 2.5. Statistical Analysis

Demographic data are presented as mean ± standard deviation. Comparisons of numbers of particles between regions and delivery type proceeded in an iterative fashion, with data for MPs and NPPs evaluated separately. Initially, a 2-way ANOVA was attempted but the data did not meet the essential assumptions of normality (Shapiro Wilk test) and equal variance (Brown-Forsythe test) required for a parametric test. Subsequently, the effects of delivery on particle numbers in each placenta region or summed across all regions within each placenta were analysed by 2-sided T-test (if normal based on Shapiro Wilk test) or Mann-Witney rank sum test. Effects of delivery method were tested using the Friedman test, in which data for particle numbers in each separate region were considered repeated measures within each sampled placenta. Differences in particle numbers between placenta region were analysed using the Kruskal Wallis test, as normality testing failed. Significance for all statistical tests was set at p<0.05. As no tests indicated significant differences, no post-hoc comparisons were used. All statistical analyses were carried out using SigmaPlot (v13.0, Systat Inc, San Jose, CA).

## 3. Results

### 3.1 Patient characteristics

Patient demographics, stratified according to mode of delivery, are presented in **Table 1**. Patients who delivered by C-section had slightly lower gestational ages compared to those delivered vaginally, but this failed to reach statistical significance (38.2 ± 1.30 vs 39.8 ± 1.09; p = 0.07). All patients who delivered by C-section were carrying male fetuses (5/5), compared to 60% of patients in the vaginal delivery group (3/5). No differences were noted for maternal age, birthweight and placental weight, and no smokers were included in either group of patients.

**Table 1.** Patient Demographics.

|  | Vaginal Delivery<br>(n=5) | C-Section Delivery<br>(n=5) | T-Test P-Value |
| --- | --- | --- | --- |
| Maternal Age; yrs | 34 ± 3.71<sup>1</sup> | 33 ± 2.55 | 0.507 |
| Smoker; n (%) | 0 | 0 | - |
| Gestational age at delivery; weeks | 39.8 ± 1.09 | 38.2 ± 1.30 | 0.07 |
| Fetal Sex; n males (% males) | 3 (60) | 5 (100) | - |
| Birthweight percentile | 46 ± 34.5 | 72.6 ± 29.7 | 0.23 |
| Placental weight; g | 663 ± 133 | 685 ± 121 | 0.79 |

<sup>1</sup> Data presented as mean + std dev.

### 3.2 Post-delivery and environmental MP contamination

Four plastic materials were identified in the delivery and surgical suites of the birthing unit that could not be substituted during the delivery and sampling protocol and underwent MP evaluation. Only one item tested, the surgical drape used during C-section deliveries, demonstrated evidence of MP shedding, with 3 MP particles identified in 30 ml of rinsate, identified as polypropylene (PP) polymers (**Supplemental Table 1**).

In both HBs and PBs some contaminating fibres were identified, with cotton or human hair source origin (**Supplemental Table 2**). Of note, these contaminating fiber types would be completely degraded in the sample digestion protocol employed. Nevertheless, fibres were excluded from all enumeration of observed particles in our placenta samples, regardless of their composition. Other types of contaminations shown in **Supplemental Table 2** were observed in the HBs from C-section sample 1 (C1), C4, Vaginal sample 1 (V1), V2 and V3 and in PBs of C3, C4, V2 and V3. However, these contaminants were infrequently observed and did not appear to be commonly found in any biological samples tested.

### 3.3 Identification of MPs in Placenta Samples

For each patient, 1 gram of tissue was collected from each of the three distinct anatomical regions of the placenta and processed for Raman Microspectroscopy (**Figure 1**). Identified MPs were noted in all placentas sampled, with a total of 31 MP particles detected across all ten placentas. The number of MPs per placenta varied (ranging from 1-11, **Figure 2**), with an average of 1 ± 1.2 MP/g of tissue. The identified MPs ranged in size from approximately 2 - 60 μm. No significant differences were observed in the number of identified MPs across the three anatomical regions sampled – the basal plate, the chorionic villous and chorionic plate (p>0.05; **Figure 2**). There was likewise no difference in the total or localized number of MPs identified according to mode of delivery (p>0.05; **Figure 2**)

**Figure 1.**
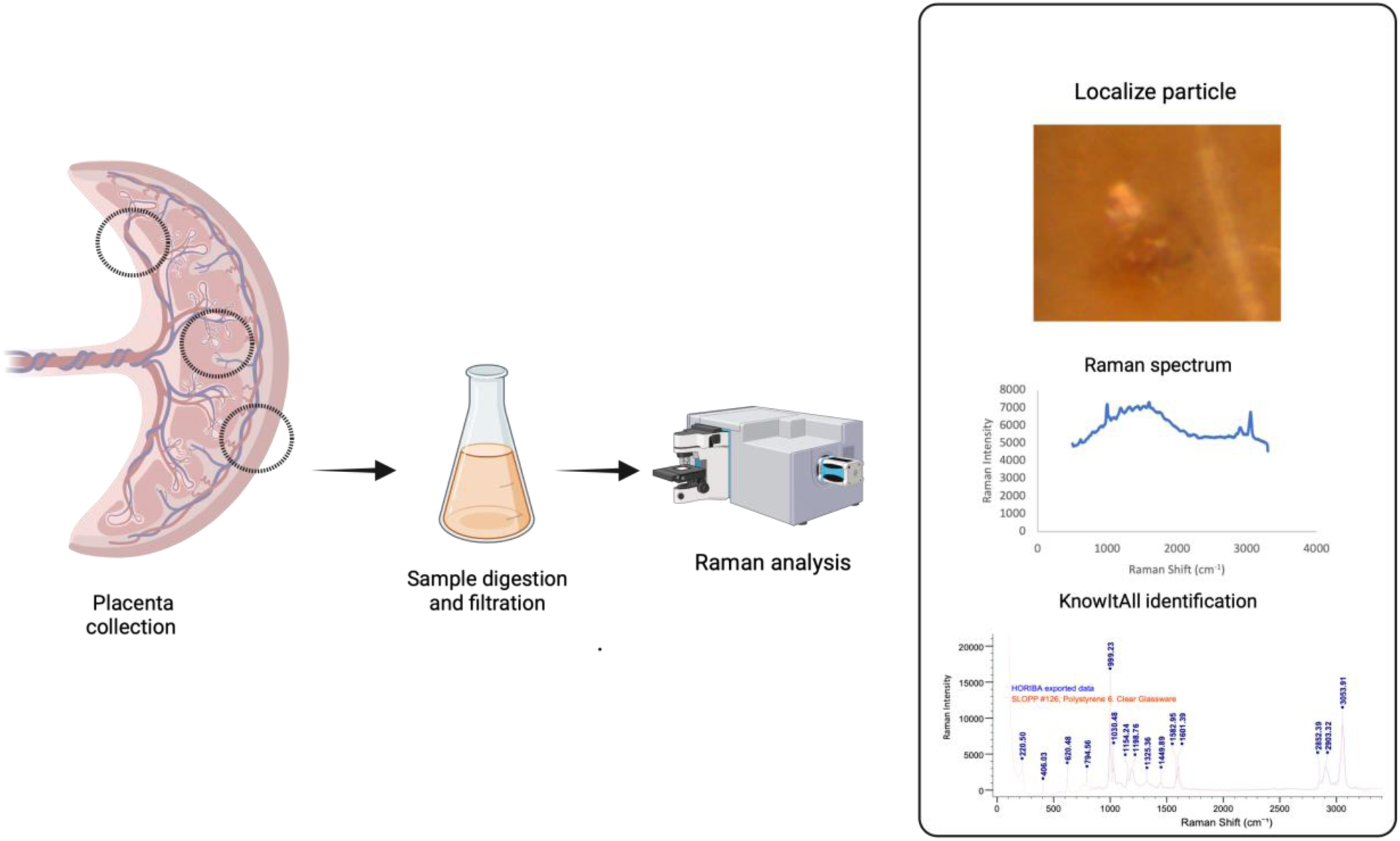
Methodology workflow of placenta tissue sampling and processing for microplastic detection. One gram of placental tissue was collected from each anatomical site (chorionic plate, chorionic villus, and basal plate), which underwent chemical digestion and filtering through a GF/D membrane. Microplastics were identified using Raman micro-spectroscopy, with the generated spectra compared to the KnowItAll database to determine the particulate identity.

**Figure 2.**
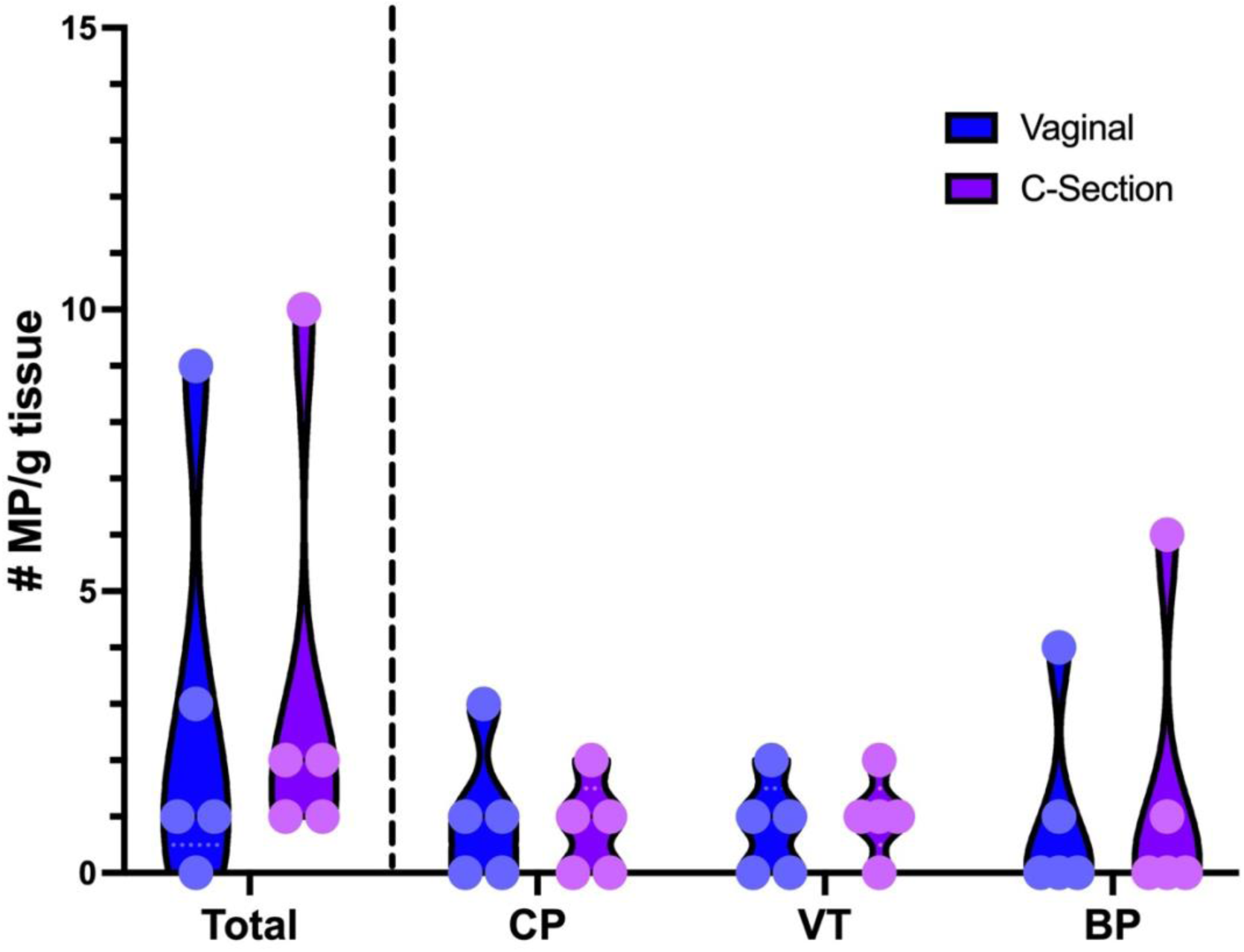
Total and region-specific number of microplastics (MP) identified in placenta tissue collected from vaginal (blue) and C-section (purple) deliveries. One gram of tissue was collected and processed from each anatomical site for each placenta. BP = basal plate; VT = chorionic villous tissue; CP = chorionic plate.

### 3.4 Characterization of MPs identified within placenta samples

Raman spectra were analyzed for each identified particle and exported into the KnowItAll database software for comparison to a library of spectra from known materials. In the collected placenta samples, the most abundant MP detected was unidentified polymers containing Copper-phthalocyanine, found in 6/10 placentas (range: 0 - 10 /g tissue), however no significant differences were observed according to anatomical location within the placenta or mode of delivery (P=0.54; **Tables 2, 3**). Polyethylene (PE), polystyrene (PS) and polyvinyl chloride (PVC) were each identified in 2/10 placentas (with 0.3 particles/g tissue for each polymer type and sample). Other common polymers identified in at least one placenta, included PP and polymethyl methacrylate (PMMA) (both with 0.3 particles/g tissue). While PE was only observed in 1/10 placentas sampled, a total of 7 particles of PE were found across all three placental compartments in that placenta (range: 2 - 3 MP/g tissue). As sample numbers were too small, statistical analysis of differences in MP polymer type according to placental distribution or mode of delivery was not possible for PP, PMMA, and PE. Representative bright field images for each of these MPs are depicted in **Figure 3**, with detailed spectra analysis for each presented in **Supplemental Table 3**. Additional polymers were observed, including 1 styrene isoprene and 1 phthalocyanine (Mortoperm blue - polymer dye), however the polymer matrix containing the dye was not identifiable.

**Figure 3.**
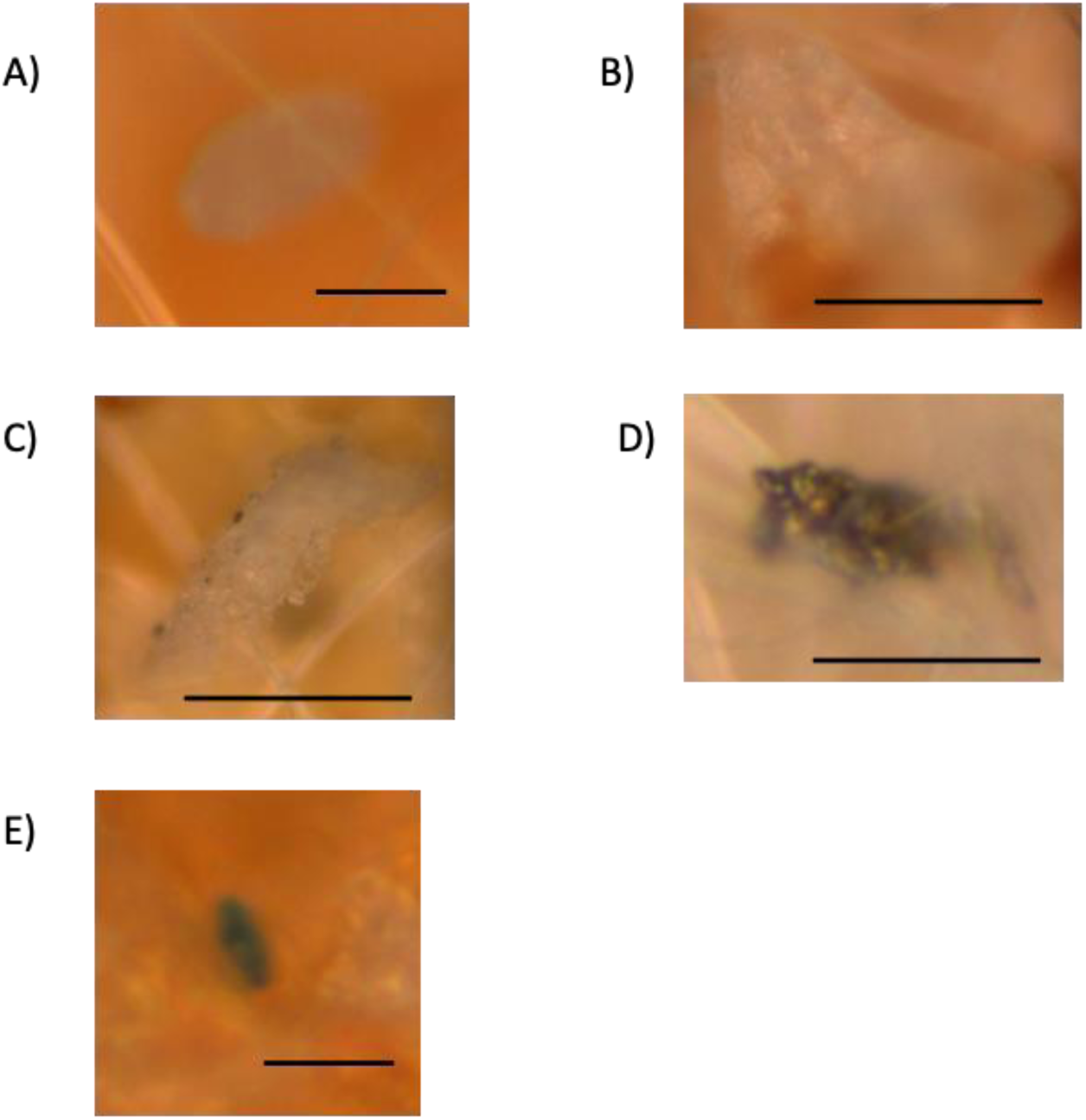
Representative brightfield images of common types of microplastic polymers found in human placenta samples. **A)** Polyethylene (PE) – scale bar represents 5µm, **B)** Polypropylene (PP) – 10µm, **C)** Polystyrene (PS) – 10µm, **D)** Polyvinyl Chloride (PVC) – scale bar 10µm, **E)** Copper phthalocyanine – 5µm.

**Table 2.** Total number, distribution, and polymer character of detected microplastic particles per gram tissue in placentas collected following C-section and vaginal deliveries.

| MP Polymer | C-section (n= 5) |  |  |  | Vaginal (n=5) |  |  |  |
| --- | --- | --- | --- | --- | --- | --- | --- | --- |
|  | BP<br>(MP/g<br>tissue) | VT<br>(MP/g<br>tissue) | CP<br>(MP/g<br>tissue) | Size<br>(um) | BP<br>(MP/g) | VT<br>(MP/g<br>tissue) | CP<br>(MP/g<br>tissue) | Size<br>(um) |
| PE | - | - | - | - | 2 | 2 | 3 | 5 – 20 |
| PP | 1 | - | - | 60 | - | - | - | - |
| PS | - | 1 | - | 7 | 1 | - | - | 40 |
| PVC | - | 2 | - | 9 - 15 | - | - | - | - |
| PMMA | - | - | 1 | 3 | - | - | - | - |
| Copper<br>phthalocyanine<sup>#</sup> | 6 | 3 | 2 | 2 – 40 | 2 | 1 | 2 | 12 - 44 |
| Styrene-isopropene | - | - | 1 | 10 | - | - | - | - |
| Mortoperm blue<sup>#</sup> | - | - | - | - | 1 | - | - | 7 |
| All polymers | 7 | 6 | 4 | 2 - 60 | 6 | 3 | 5 | 5 - 44 |

<sup>#</sup>Dye used in plastic.
MP = microplastic; g = gram tissue; BP = Basal Plate; VT = Chorionic Villous Tissue; CP =
Chorionic Plate; PE = Polyethylene; PS = Polystyrene; PP = Polypropylene; PVC = Polyvinyl
Chloride; PMMA = Polymethyl methacrylate.

**Table 3.** The number of microplastic particles identified in placenta tissue according to polymer type and mode of delivery.

| MP Polymer | C-Section (n= 5)<br># MPs identified* |  |  |  |  | Vaginal delivery (n = 5)<br># MPs identified* |  |  |  |  | Total # MPs<br>identified by<br>polymer type |
| --- | --- | --- | --- | --- | --- | --- | --- | --- | --- | --- | --- |
|  | C1 | C2 | C3 | C4 | C5 | V1 | V2 | V3 | V4 | V5 |  |
| PE | - | - | - | - | - | - | - | 7 | - | - | 7 |
| PP | - | - | - | 1 | - | - | - | - | - | - | 1 |
| PS | 1 | - | - | - | - | - | 1 | - | - | - | 2 |
| PVC | - | - | 1 | 1 | - | - | - | - | - | - | 2 |
| PMMA | - | - | 1 | - | - | - | - | - | - | - | 1 |
| Copper<br>phthalocyanine<sup>#</sup> | - | 1 | - | - | 10 | 1 | 1 | 2 | - | 1 | 16 |
| Styrene-isopropene | - | - | - | - | 1 | - | - | - | - | - | 1 |
| Mortoperm blue<sup>#</sup> | - | - | - | - | - | - | - | - | 1 | - | 1 |
| **Total # MPs**<br>**identified/**<br>**Placenta\*** | **1** | **1** | **2** | **2** | **11** | **1** | **2** | **9** | **1** | **1** | **31** |

\*Total of 3 grams collected per placenta.
<sup>#</sup>Dye used in some plastic.
Polyethylene (PE); Polystyrene (PS); Polypropylene (PP); Polyvinyl Chloride (PVC); Polymethyl methacrylate (PMMA).

### 3.5 Identification of non-MPs particles in placenta samples

Raman spectroscopy revealed that most particles observed in digested placenta samples were not MPs: All placenta samples contained NPPs, with a total of 121 NPPs detected across all 10 placentas. The number of NPPs per placenta varied (ranging from 2-34, **Figure 4**), with an average of 4 ± 2.9 NPPs/g of tissue. The identified NPPs ranged in size from 2 - 100 μm (**Table 4**). There was no significant difference in the total number of NPPs identified according to mode of delivery (p= 0.155; **Figure 4**, **Table 4**).

**Fig 4.**
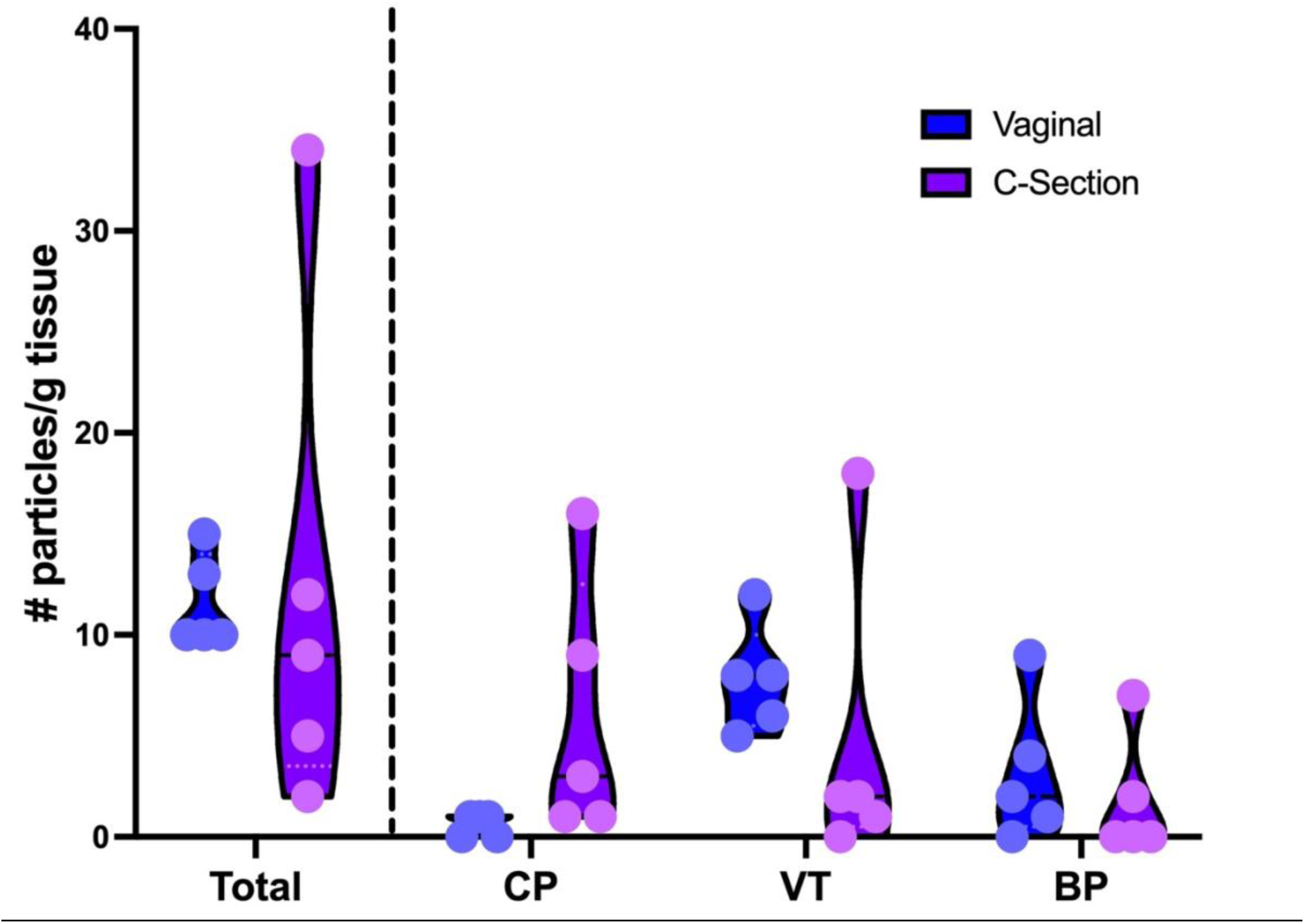
Total and region-specific number of non-microplastic particles identified in placenta tissue collected from vaginal (blue) and C-section (purple) deliveries. One gram of tissue was collected and processed from each anatomical site for each placenta. BP = basal plate; VT = chorionic villous tissue; CP = chorionic plate.

**Table 4.** Total number, distribution, and polymer character of detected non-MP particles per gram tissue in placentas collected following C-section and vaginal deliveries.

| Non-MP<br>Particle | C-section (n= 5) |  |  |  | Vaginal (n=5) |  |  |  |
| --- | --- | --- | --- | --- | --- | --- | --- | --- |
|  | BP<br>(particles<br>/g tissue) | VT<br>(particles<br>/g tissue) | CP<br>(particles<br>/g tissue) | Size (um) | BP<br>(particle<br>s/g<br>tissue) | VT<br>(particle<br>s/g<br>tissue) | CP<br>(particle<br>s/g<br>tissue) | Size<br>(um) |
| Carbon<sup>#</sup> | 5 | - | - | 10 - 25 | - | 3 | 2 | 3 - 15 |
| Graphite | 5 | 4 | - | 2 - 25 | - | 8 | 1 | 5 - 16 |
| Lead<br>Oxide | 4 | 5 | 1 | 12 - 40 | - | - | - | - |
| Bayerite | 3 | 2 | - | 6 - 60 | - | 5 | - | 12 - 34 |
| Other<sup>^</sup> | 11 | 10 | 3 | 3 - 80 | 3 | 23 | 11 | 4 - 100 |
| Unknown<sup>+</sup> | 2 | 6 | 1 | 3 - 40 | - | 1 | 2 | 8 - 80 |

<sup>#</sup> Includes black carbon, diamond like carbon
<sup>^</sup>Includes less frequently found polymers, details included in **Supplemental Table 4**
<sup>+</sup>Spectrum could not be identified through Raman spectra and KnowItAll database.
Basal Plate (BP); Chorionic Villous Tissue (VT); Chorionic Plate (CP)

### 3.6 Characterization of non-MPs particles in placenta samples

Non-MP particle composition was also identified in placenta samples using Raman micro-spectroscopy, with confirmation of particle identity though KnowItAll database. A total of 29 unique particle identities were captured **(Supplemental Table 3**), with the most abundant particles identified as carbon, graphite, lead oxide and bayerite composition (**Tables 4** and **5**). Each of these particles were observed in 2-3 different placentas, ranging from 1-22 total particles per placenta (1.5 ± 1.1 particles/g of tissue). No significant differences were observed in the specific non-MP particle type according to anatomical location within the placenta or mode of delivery (P> 0.05; **Tables 4** and **5**). Representative bright field images for each of these MPs are depicted in **Figure 5**, with detailed spectra analysis for each presented in **Supplemental Table 5**. It should be noted that there were 12 particles for which Raman spectral analysis was inconclusive and, therefore, could not be identified (**Table 5**).

**Figure 5.**
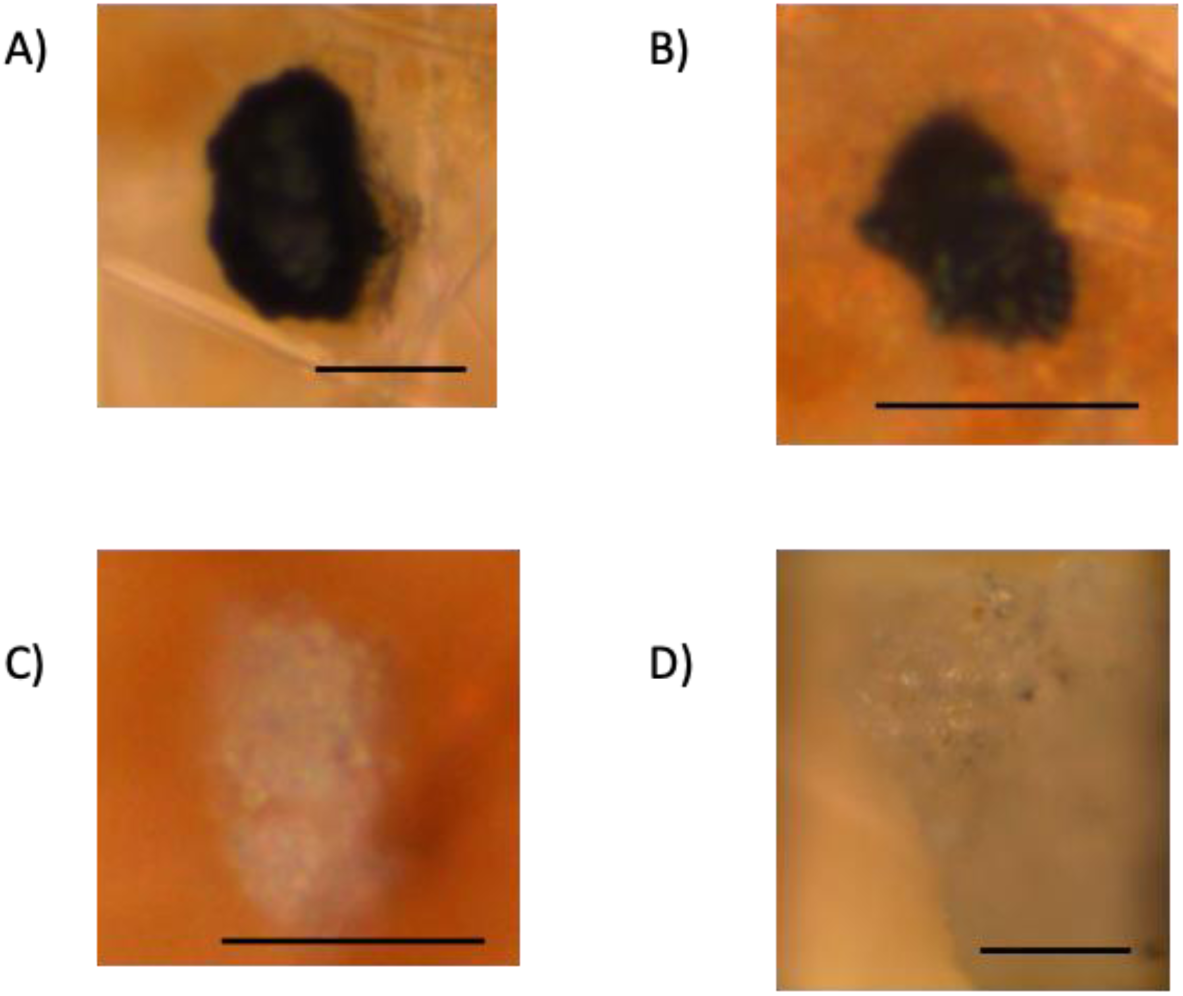
Representative brightfield images of common types of non-microplastic particles found in human placenta samples. **A)** Carbon – scale bar represents 5µm B) Graphite-10µm, **C)** Lead Oxide – 10µm, **D)** Bayerite - 5µm.

**Table 5.** The number of non-microplastic particles identified in placenta tissue according to particle type and mode of delivery.

| Particle type | C-Section (n= 5)<br># particles identified* |  |  |  |  | Vaginal delivery (n = 5)<br># particles identified* |  |  |  |  | Total # of particles<br>identified by type |
| --- | --- | --- | --- | --- | --- | --- | --- | --- | --- | --- | --- |
|  | C1 | C2 | C3 | C4 | C5 | V1 | V2 | V3 | V4 | V5 |  |
| Carbon<sup>#</sup> | 5 | - | - | - | - | 2 | - | - | 3 | - | 10 |
| Graphite | - | - | 9 | - | - | - | - | 8 | - | 1 | 18 |
| Lead Oxide | - | - | 9 | 1 | - | - | - | - | - | - | 10 |
| Bayerite | - | - | 4 | - | 1 | - | - | - | 5 | - | 10 |
| Other<sup>^</sup> | 4 | - | 6 | 11 | 3 | 11 | 13 | 2 | 2 | 9 | 61 |
| Unknown<sup>+</sup> | - | 2 | 6 | - | 1 | 2 | 1 | - | - | - | 12 |
| **Total # particles**<br>**identified/**<br>**Placenta\*** | **9** | **2** | **34** | **12** | **5** | **15** | **14** | **10** | **10** | **10** | **121** |

\*Total of 3 grams collected per placenta.
<sup>#</sup> Includes black carbon, diamond like carbon
<sup>^</sup>Includes frequently observed particle types, details included in Supplemental Table 4
<sup>+</sup>Spectrum could not be identified through Raman spectra and KnowItAll database.

## 4. Discussion

The major findings of this study indicate the presence of both MP and NPPs in all placentas sampled, distributed across all placental compartments and unrelated to the mode of delivery. This small sample size was selected from individuals whose singleton pregnancies were uncomplicated, delivering within a tertiary care hospital setting, in a large Canadian city (Ottawa, Ontario). While these exposure levels may not represent the diversity of exposures across the Canadian population, it provides insight regarding routine particulate exposures of pregnant individuals within a major urban region in Canada. To the best of our knowledge this study is the first to report presence of MPs in human placentas within a Canadian population.

Our findings are mostly consistent with those reported in recent publications (Amereh et al., 2022; Braun et al., 2021; Garcia et al., 2024; Ragusa et al., 2021; Weingrill et al., 2023). Importantly, the current study found MP contamination in all placentas sampled (100%, n=10), whereas smaller observational studies carried out in Italian and German urban city obstetrical populations (Braun et al., 2021; Ragusa et al., 2021) have reported MPs in less than 70% of placentas analysed. The current study further identified the presence of NPPs in all placentas sampled, many composed of synthetic materials. Few studies have reported the presence of NPPs in human placenta tissues to date, although Ragusa et al., did describe the presence of unidentified particles bound to polymer dyes (Ragusa et al., 2022a, 2021). Carbon was also a common NPP placental contaminant identified in the current study, findings that have been previously described by others (Bové et al., 2019) and found to correlate with levels of black carbon found in dust particulate within an individual’s home during pregnancy.

An important consideration when interpreting the scope of the presented MP (1 ± 1.2 MPs/g tissue) and NPPs (4 ± 2.9 non-MPs/g tissue) accumulation within the placenta, is the total size of this organ of pregnancy at the time of delivery – typically ranging from 550-650 grams. As such, in our sampled Canadian population, we estimate that the average placenta may contain upwards of 650 MPs and 2,600 NPPs at the end of gestation. A second critical consideration for data interpretation, is the current technical limitations of MP and NPPs identification by Raman Micro-spectroscopy at the lower end of the particle size spectrum (<1 μm). The smallest particles detected in the current study were ∼2 µm – aligned with both the pore size of the glass fibre filters used to harvest particles from the sample digests and the lower threshold of detection of this current technology (Araujo et al., 2018). Due to the higher bioavailability of smaller particles throughout the human body and placenta (Wick et al., 2010), it is likely that smaller particles are present in the placenta that were not accounted for in the presented measurements. As such, the reported concentrations are underestimates of total particle accumulation in the placenta. Importantly, we did note considerable variability in total placenta particle load between individuals, ranging from 0.3 to 11.3 particles/g tissue. No smokers were included within the sampled population; however, these results would indicate vastly different particle exposure levels during pregnancy, possibly related to differences in diet/food practices, socioeconomic status, workplace, home life and other environmental exposures. The placenta is also a large and highly heterogenous organ, and as such there may also be variability based on sampling site. Future work, focused on better understanding the associations between gestational MP/NPPs exposures and geo-demographical variables, along with placental distribution is warranted.

The chemical character of MP and NPPs were identified for most particles detected in the current study – with the most common MPs identified as PE, PP, PS and PVC, and the most common NPP identified as carbon, graphite and lead oxide. While these characterizations are important for understanding relevant gestational exposures, concrete determination of original sources of the exposures is not possible. However, we can look to commonly known sources to gain perspective on potential sources. PE is widely used in packaging – including food packaging - and items with short functional lifespans including plastic bags, plastic cling film packaging, bottles, etc (Geyer et al., 2017; Kadac-Czapska et al., 2023). PP is widely used in packaging, transportation of materials, and consumer products including plastic bottles, indoor-outdoor carpets, microwavable containers, and disposable face masks (Geyer et al., 2017; “Science of Plastics,” n.d.). PS is found as a solid or expanded foam (styrofoam) and has a broad array of uses including many of single use products used in biomedical applications, disposable food containers and cutlery and packaging (American Chemistry Council, 2022; Geyer et al., 2017) while PVC has diverse uses in building material, consumer products, be found in plastic pipes, synthetic leather, clear food wrap, flooring materials and soft toys, among others (Geyer et al., 2017). Likewise, the most abundant NPPs identified are pervasive in the environment. Black carbon particle is a major constituent of air pollution, the result of incomplete combustion of fossil fuels (“What is Black Carbon?,” n.d.) .There are multiple industrial uses of natural and synthetic graphite that result in significant environmental release of airborne particulates (Zhang et al., 2023), whereas lead oxide particles could arise from the use of lead in old batteries, from paint pigments (especially white), or gas sensors (Bratovcic, 2020). The frequency of use of these products and materials represents a wide range of potential exposures to humans.

The distribution of MPs and NPPs across all three regions of the placenta sampled suggest that the placenta does not act as a barrier to fetal particulate exposure. Had there been a notable concentration uniquely located on the maternal surface (basal plate), one could have inferred a selective barrier function of the placenta regarding MP and NPPs transfer to the fetal compartment. The presence of MPs throughout the placenta certainly warrants concern regarding the functional impact of these particles on placental integrity and function (i.e., maternal-fetal exchange, hormone production etc.) and certainly infer MP translocation into the fetal compartment. Indeed, ex vivo placenta perfusion studies have confirmed the transfer of PS nanoplastics from maternal to fetal compartments (Grafmueller et al., 2015; Wick et al., 2010). Furthermore, the detection of MPs in the meconium of newborn infants (Braun et al., 2021; Liu et al., 2022a, 2022b) and amniotic fluid at term (Xue et al., 2024) provides clear evidence of in utero fetal exposure to MPs.

The widespread distribution of MPs across all placenta samples underscores the significance of investigating their potential consequences on placental function and fetal development, an area of investigation that has been relatively unexplored, particularly in human populations. Evidence collected using rodent models of gestational MP exposure have demonstrated smaller placentas (Fournier et al., 2020; Hu et al., 2021; Nie et al., 2021), impaired feto-placental and utero-placental vasculature development (Chen et al., 2022; Hu et al., 2021), disturbances in uterine immune cell balance (Hu et al., 2021) and placental metabolic disruption (Aghaei et al., 2022; Chen et al., 2022) . In vitro exposure of human placental tissue to MPs has also been shown to alter expression of genes and proteins related to inflammation and iron homeostasis (Chortarea et al., 2023). A recent study showed an inverse correlation between the accumulation of MPs in the placenta and birthweight (correlation coefficient, r = -0.82, p < 0.001), with similar associations observed for neonatal length at birth, head circumference, and 1-minute APGAR scores (Amereh et al., 2022). . In addition, the levels of MPs detected in amniotic fluid collected at delivery was inversely related to gestational age (Xue et al., 2024) suggesting that in utero exposures to MPs tends to cause earlier labour. Notably, non-MP black carbon particles have also been identified in higher concentrations within placentas of pregnancies complicated by pre-term delivery and/or FGR, compared to health controls (Amereh et al., 2022; Bové et al., 2019).

Analysis was also conducted to assess the impact of the mode of delivery, specifically caesarean and vaginal deliveries, on the presence of both MPs and non-MP particles within placental tissues. The objective was to discern whether any disparities in the quantity or types of MPs detected in placentas from these delivery modes could shed light on whether contamination occurred post-delivery or resulted from in utero accumulation during placental development. The absence of significant variations in MPs and non-MPs levels between the two delivery modes supports the hypothesis that MPs contaminate the placental tissue during gestation rather than after birth. This outcome also reflects the effectiveness of the measures taken to minimize plastic contamination throughout the delivery process, placenta dissection, sample collection, storage, digestion, and all other phases of sample handling and analysis. This confidence in the prevention of contamination is further supported by the near absence of particle contamination of PBs, which served to monitor potential sources of contamination during sample dissection and sample processing. In instances where particles were noted in the PBs, they did not bear resemblance to the particles found within the placenta samples. Furthermore, the presence of particles within the chorionic villous region of the placenta, located deep within the organ and shielded from external contact during delivery, provides evidence that these particles were present *in utero*. These collective observations strengthen the conclusion that MP particles identified in placental samples are indicative of in utero exposure, rather than due to post-delivery contamination, and contribute to the growing body of evidence that particulate pollution reaches into the human womb.

## 5. Conclusions

This study observed both MPs and non-plastic foreign particles in human placentas collected at term. Particles were found in all placentas tested, and throughout the various compartments of the placenta – demonstrating the ability of these particles to translocate into the fetal compartment. As the placenta is the vital organ of pregnancy supporting fetal development, any adverse effects of these foreign particles on the health and function of the placenta can have serious effects on the health and wellbeing of the offspring. Further research is needed to understand the repercussion of this exposure in pregnancy on the gestational parent and the developing fetus.

## Supplement

**Supplemental Table 1.** MP shedding analysis of plastic materials used in birthing suites at The Ottawa Hospital that could not be replaced in the plastic-reduced protocol.

| Item | Common MPs |  |  |
| --- | --- | --- | --- |
|  | PE | PS | PP |
| Surgical drape | - | - | 3 |
| Placenta collection bowl | - | - |  |
| Placenta collection bag | - | - | - |
| Gloves | - | - | - |

**Supplemental Table 2.**
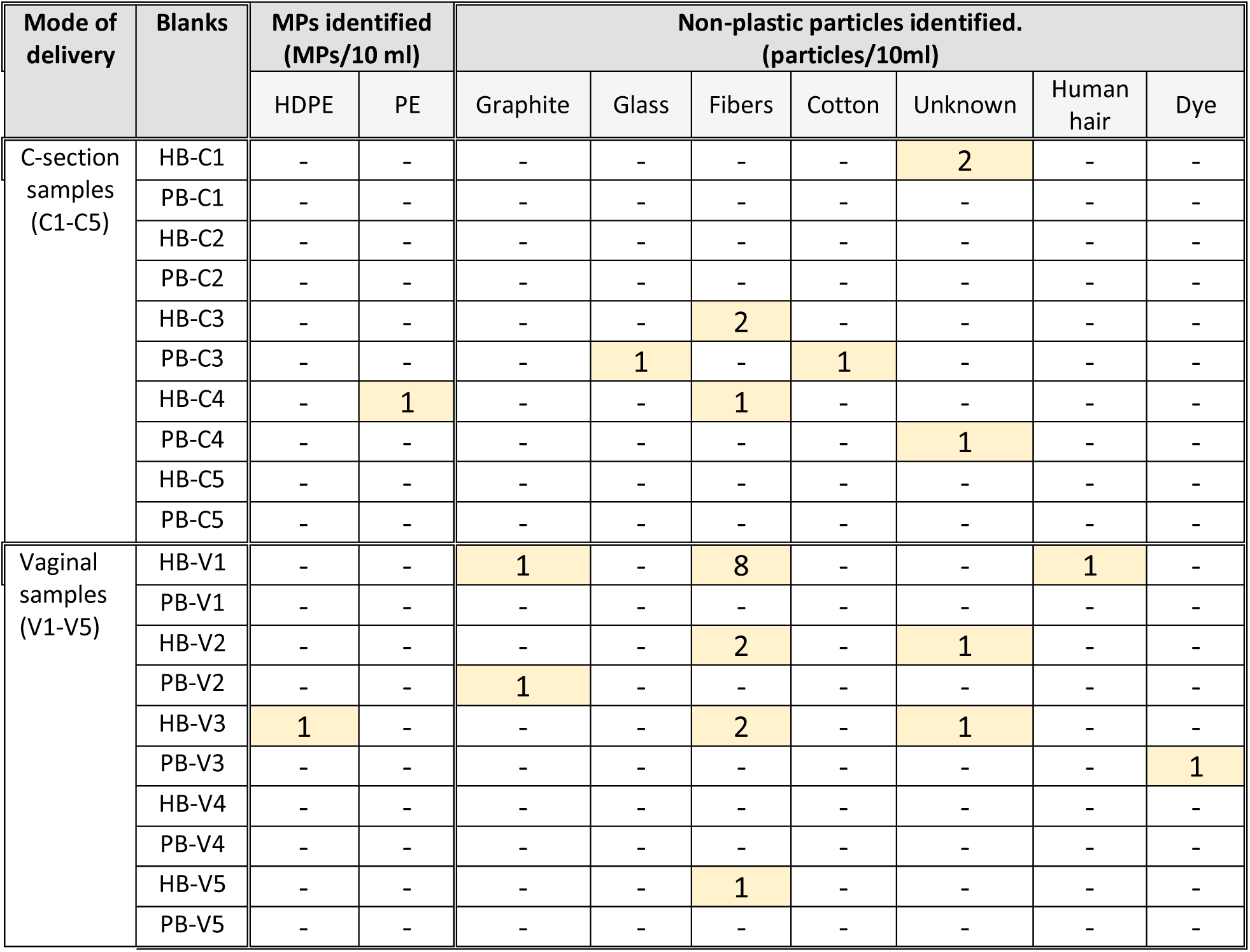
Microplastic and non-microplastic particle contamination identified in hood blanks (HB) and procedural blanks (PB) for each placenta sampling.

**Supplemental Table 3.**
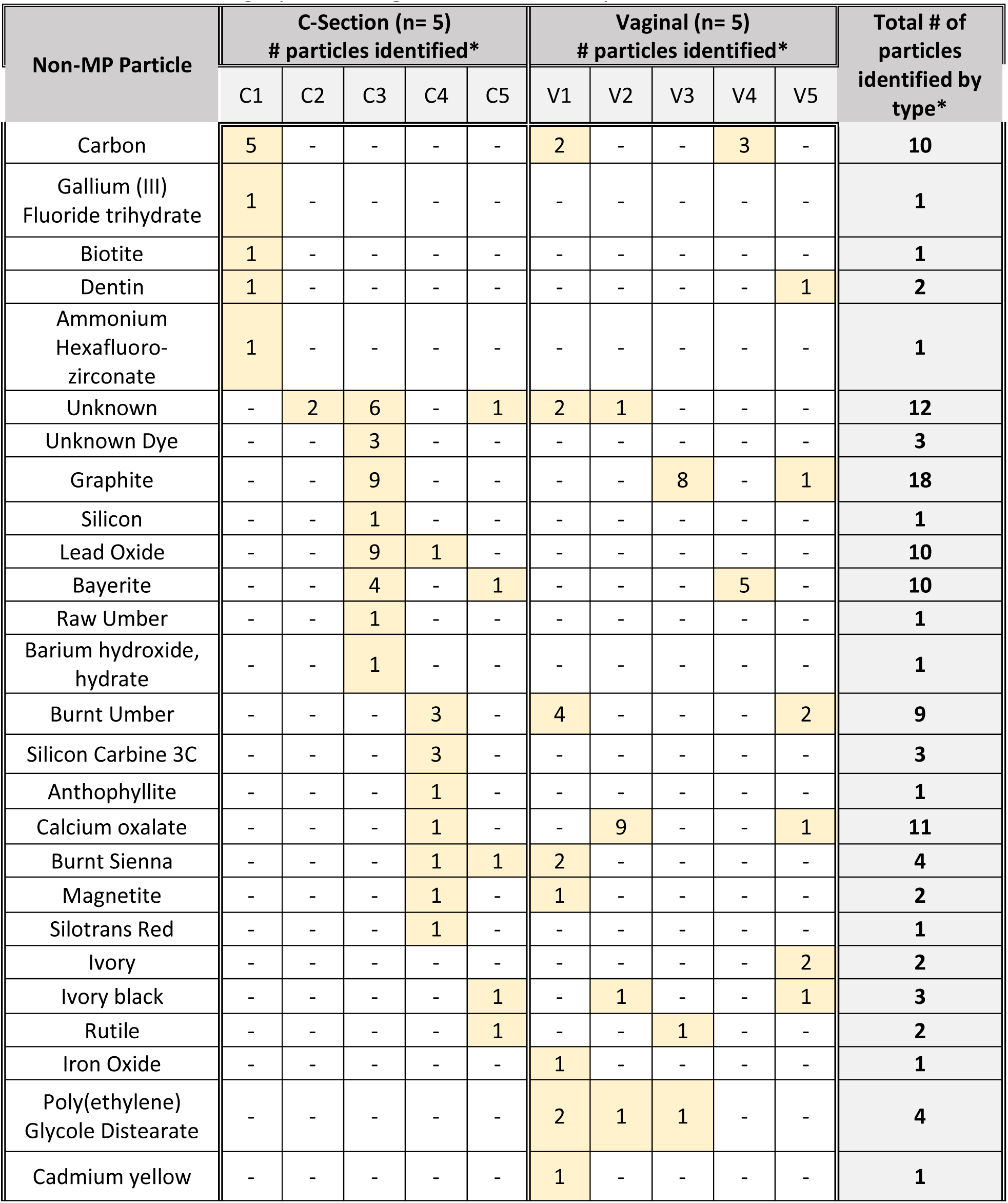
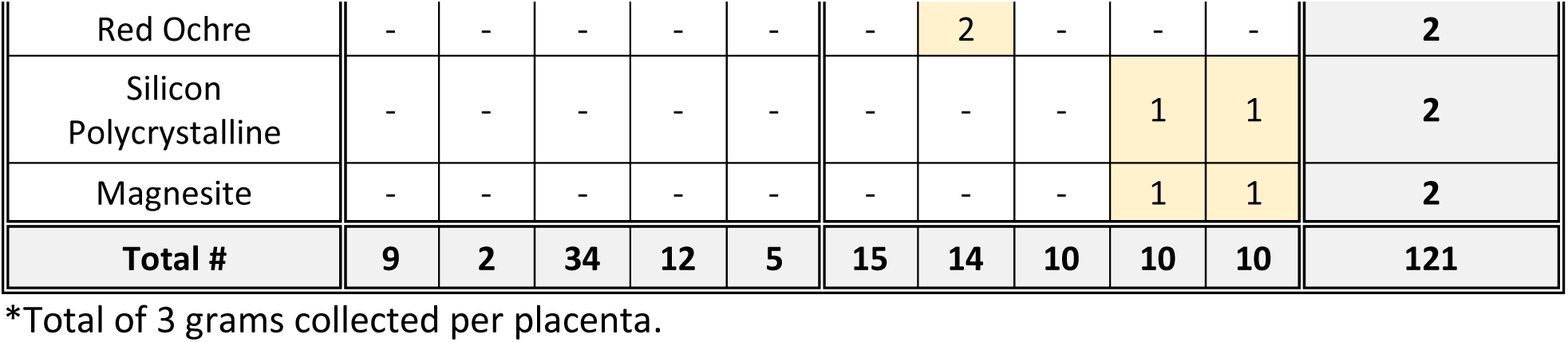
Non-MP particles identified in placenta samples that were classified within the “other” category, according to mode of delivery.

| Non-MP Particle | C-Section (n= 5)<br># particles identified* |  |  |  |  | Vaginal (n= 5)<br># particles identified* |  |  |  |  | Total # of<br>particles<br>identified by<br>type* |
| --- | --- | --- | --- | --- | --- | --- | --- | --- | --- | --- | --- |
|  | C1 | C2 | C3 | C4 | C5 | V1 | V2 | V3 | V4 | V5 |  |
| Carbon | 5 | - | - | - | - | 2 | - | - | 3 | - | 10 |
| Gallium (III)<br>Fluoride trihydrate | 1 | - | - | - | - | - | - | - | - | - | 1 |
| Biotite | 1 | - | - | - | - | - | - | - | - | - | 1 |
| Dentin | 1 | - | - | - | - | - | - | - | - | 1 | 2 |
| Ammonium<br>Hexafluoro-<br>zirconate | 1 | - | - | - | - | - | - | - | - | - | 1 |
| Unknown | - | 2 | 6 | - | 1 | 2 | 1 | - | - | - | 12 |
| Unknown Dye | - | - | 3 | - | - | - | - | - | - | - | 3 |
| Graphite | - | - | 9 | - | - | - | - | 8 | - | 1 | 18 |
| Silicon | - | - | 1 | - | - | - | - | - | - | - | 1 |
| Lead Oxide | - | - | 9 | 1 | - | - | - | - | - | - | 10 |
| Bayerite | - | - | 4 | - | 1 | - | - | - | 5 | - | 10 |
| Raw Umber | - | - | 1 | - | - | - | - | - | - | - | 1 |
| Barium hydroxide,<br>hydrate | - | - | 1 | - | - | - | - | - | - | - | 1 |
| Burnt Umber | - | - | - | 3 | - | 4 | - | - | - | 2 | 9 |
| Silicon Carbide 3C | - | - | - | 3 | - | - | - | - | - | - | 3 |
| Anthophyllite | - | - | - | 1 | - | - | - | - | - | - | 1 |
| Calcium oxalate | - | - | - | 1 | - | - | 9 | - | - | 1 | 11 |
| Burnt Sienna | - | - | - | 1 | 1 | 2 | - | - | - | - | 4 |
| Magnetite | - | - | - | 1 | - | 1 | - | - | - | - | 2 |
| Silotrans Red | - | - | - | 1 | - | - | - | - | - | - | 1 |
| Ivory | - | - | - | - | - | - | - | - | - | 2 | 2 |
| Ivory black | - | - | - | - | 1 | - | 1 | - | - | 1 | 3 |
| Rutile | - | - | - | - | 1 | - | - | 1 | - | - | 2 |
| Iron Oxide | - | - | - | - | - | 1 | - | - | - | - | 1 |
| Poly(ethylene)<br>Glycole Distearate | - | - | - | - | - | 2 | 1 | 1 | - | - | 4 |
| Cadmium yellow | - | - | - | - | - | 1 | - | - | - | - | 1 |
| Red Ochre | - | - | - | - | - | - | 2 | - | - | - | 2 |
| Silicon Polycrystalline | - | - | - | - | - | - | - | - | 1 | 1 | 2 |
| Magnesite | - | - | - | - | - | - | - | - | 1 | 1 | 2 |
| **Total #** | **9** | **2** | **34** | **12** | **5** | **15** | **14** | **10** | **10** | **10** | **121** |

\*Total of 3 grams collected per placenta.

**Supplemental Table 4.** Non-plastic particles and potential sources.

| **Particles** | **Potential sources** |
| --- | --- |
| **Carbon** | Product of air pollution due to incomplete combustion of fossil and other fuels, photo copiers, printers, coloring agent. |
| **Gallium (III) Fluoride trihydrate** | water gallium source for use in oxygen-sensitive applications such as metal production |
| **Biotite** | Silicate mineral in the common mica group. Found in granite (counter tops) |
| **Dentin** | A thick dentin layer forms the bulk of dental mineralized dental tissue |
| **Ammonium Hexafluorozirconate** | chemical compound used in corrosion-resistant adherent coatings on aluminum surfaces and aqueous acid solutions for chemical conversion coatings |
| **Graphite** | Crystalline form of the element carbon; Found in pencils, lubricants, lamps, batteries, polishes |
| **Silicon** | Used in cookware, bakeware, toys, adhesive, lubricants, gaskets, medical application |
| **Lead Oxide** | Widely used in batteries, gas sensors, pigments and paints, ceramics, glass |
| **Bayerite** | Production of aluminum metals and used in flame retardants |
| **Raw Umber** | Umber is a natural brown earth pigment color that contains iron oxide and manganese oxide. In its natural form it is called raw umber |
| **Barium hydroxide, hydrate** | Highly water insoluble crystalline barium source for uses compatible with higher basic pH environments |
| **Burnt Umber** | Umber is a natural brown earth pigment, made by heating raw umber, which dehydrates the iron oxide |
| **Silicon Carbide 3C** | Silicon Carbide 3C is found in ceramic plates, bulletproof vest, car clutches. Silicon carbide is a hard chemical compound containing silicon and carbon |
| **Anthophyllite** | Type of asbestos |
| **Calcium oxalate** | Oxalate is found in certain foods such as spinach, almonds, soy etc. Oxalate cannot be metabolized and must be excreted through urine. Elevated levels may lead to kidney stones |
| **Burnt Sienna** | Earth pigment from iron oxide |
| **Magnetite** | A mineral whose primary component is iron oxide; Greatest use is iron ore for steel manufacture |
| **Silotrans Red** | Dye - not sure for what specifically |
| **Ivory** | Piano keys, jewelry, chess sets, also used in medicines |
| **Ivory black** | Black paint by carbonising ivory |
| **Rutile** | Oxide mineral composed titanium dioxide |
| **Iron Oxide** | Chemical composed of iron and oxygen |
| **Poly(ethylene) Glycole Distearate** | cosmetic products as surfactants, emulsifiers, cleaning agents and skin conditioners |
| **Cadmium yellow** | Yellow pigment/paint |
| **Red Ochre** | A variant of ochre containing a large amount of hematite, or dehydrated iron oxide, has a reddish tint known as red ochre |
| **Silicon Polycrystalline** | unsure |
| **Magnesite** | a refractory material, a catalyst and filler in the production of synthetic rubber, and a material in the preparation of magnesium chemicals and fertilizers. |

## Reference

Afrin, S., Rahman, M.M., Akbor, M.A., Siddique, M.A.B., Uddin, M.K., Malafaia, G., 2022. Is there tea complemented with the appealing flavor of microplastics? A pioneering study on plastic pollution in commercially available tea bags in Bangladesh. Sci. Total Environ. 837, 155833. 10.1016/j.scitotenv.2022.155833

Aghaei, Z., Mercer, G.V., Schneider, C.M., Sled, J.G., Macgowan, C.K., Baschat, A.A., Kingdom, J.C., Helm, P.A., Simpson, A.J., Simpson, M.J., Jobst, K.J., Cahill, L.S., 2022. Maternal exposure to polystyrene microplastics alters placental metabolism in mice. Metabolomics Off. J. Metabolomic Soc. 19, 1. 10.1007/s11306-022-01967-8

Amato-Lourenço, L.F., Carvalho-Oliveira, R., Júnior, G.R., Dos Santos Galvão, L., Ando, R.A., Mauad, T., 2021. Presence of airborne microplastics in human lung tissue. J. Hazard. Mater. 416, 126124. 10.1016/j.jhazmat.2021.126124

Amereh, F., Amjadi, N., Mohseni-Bandpei, A., Isazadeh, S., Mehrabi, Y., Eslami, A., Naeiji, Z., Rafiee, M., 2022. Placental plastics in young women from general population correlate with reduced foetal growth in IUGR pregnancies. Environ. Pollut. 1987 314, 120174. 10.1016/j.envpol.2022.120174

American Chemistry Council, 2022. Polystyrene [WWW Document]. Chem. Saf. Facts. URL http://www.chemicalsafetyfacts.org/chemicals/polystyrene/ (accessed 2.23.24).

Andrady, A.L., 2017. The plastic in microplastics: A review. Mar. Pollut. Bull. 119, 12–22. 10.1016/j.marpolbul.2017.01.082

Araujo, C.F., Nolasco, M.M., Ribeiro, A.M.P., Ribeiro-Claro, P.J.A., 2018. Identification of microplastics using Raman spectroscopy: Latest developments and future prospects. Water Res. 142, 426–440. 10.1016/j.watres.2018.05.060

Baeza-Martínez, C., Olmos, S., González-Pleiter, M., López-Castellanos, J., García-Pachón, E., Masiá-Canuto, M., Hernández-Blasco, L., Bayo, J., 2022. First evidence of microplastics isolated in European citizens’ lower airway. J. Hazard. Mater. 438, 129439. 10.1016/j.jhazmat.2022.129439

Bongaerts, E., Lecante, L.L., Bové, H., Roeffaers, M.B., Ameloot, M., Fowler, P.A., Nawrot, T.S., 2022. Maternal exposure to ambient black carbon particles and their presence in maternal and fetal circulation and organs: an analysis of two independent population-based observational studies. Lancet Planet. Health 6, e804–e811.

Bongaerts, E., Mamia, K., Rooda, I., Björvang, R.D., Papaikonomou, K., Gidlöf, S.B., Olofsson, J.I., Ameloot, M., Alfaro-Moreno, E., Nawrot, T.S., Damdimopoulou, P., 2023. Ambient black carbon particles in human ovarian tissue and follicular fluid. Environ. Int. 179, 108141. 10.1016/j.envint.2023.108141

Bové, H., Bongaerts, E., Slenders, E., Bijnens, E.M., Saenen, N.D., Gyselaers, W., Van Eyken, P., Plusquin, M., Roeffaers, M.B., Ameloot, M., 2019. Ambient black carbon particles reach the fetal side of human placenta. Nat. Commun. 10, 3866.

Bratovcic, A., 2020. Synthesis, Characterization, Applications, and Toxicity of Lead Oxide Nanoparticles, in: Lead Chemistry. IntechOpen. 10.5772/intechopen.91362

Braun, T., Ehrlich, L., Henrich, W., Koeppel, S., Lomako, I., Schwabl, P., Liebmann, B., 2021. Detection of Microplastic in Human Placenta and Meconium in a Clinical Setting. Pharmaceutics 13, 921. 10.3390/pharmaceutics13070921

Buzea, C., Pacheco, I.I., Robbie, K., 2007. Nanomaterials and nanoparticles: sources and toxicity. Biointerphases 2, MR17-71. 10.1116/1.2815690

Calderón-Garcidueñas, L., González-Maciel, A., Mukherjee, P.S., Reynoso-Robles, R., Pérez-Guillé, B., Gayosso-Chávez, C., Torres-Jardón, R., Cross, J.V., Ahmed, I.A.M., Karloukovski, V.V., Maher, B.A., 2019a. Combustion- and friction-derived magnetic air pollution nanoparticles in human hearts. Environ. Res. 176, 108567. 10.1016/j.envres.2019.108567

Calderón-Garcidueñas, L., Reynoso-Robles, R., González-Maciel, A., 2019b. Combustion and friction-derived nanoparticles and industrial-sourced nanoparticles: The culprit of Alzheimer and Parkinson’s diseases. Environ. Res. 176, 108574. 10.1016/j.envres.2019.108574

Campion, A., Smith, K.J., Fedulov, A.V., Gregory, D.Z., Fan, Y., Godleski, J.J., 2018. Identification of Foreign Particles in Human Tissues Using Raman Microscopy. Anal. Chem. 90, 8362–8369. 10.1021/acs.analchem.8b00271

Chen, G., Xiong, S., Jing, Q., van Gestel, C.A.M., van Straalen, N.M., Roelofs, D., Sun, L., Qiu, H., 2022. Maternal exposure to polystyrene nanoparticles retarded fetal growth and triggered metabolic disorders of placenta and fetus in mice. Sci. Total Environ. 854, 158666. 10.1016/j.scitotenv.2022.158666

Chortarea, S., Gupta, G., Saarimäki, L.A., Netkueakul, W., Manser, P., Aengenheister, L., Wichser, A., Fortino, V., Wick, P., Greco, D., Buerki-Thurnherr, T., 2023. Transcriptomic profiling reveals differential cellular response to copper oxide nanoparticles and polystyrene nanoplastics in perfused human placenta. Environ. Int. 177, 108015. 10.1016/j.envint.2023.108015

Diaz-Basantes, M.F., Conesa, J.A., Fullana, A., 2020. Microplastics in Honey, Beer, Milk and Refreshments in Ecuador as Emerging Contaminants. Sustainability 12, 5514. 10.3390/su12145514

Eriksen, M., Mason, S., Wilson, S., Box, C., Zellers, A., Edwards, W., Farley, H., Amato, S., 2013. Microplastic pollution in the surface waters of the Laurentian Great Lakes. Mar. Pollut. Bull. 77, 177–182. 10.1016/j.marpolbul.2013.10.007

Fournier, S.B., D’Errico, J.N., Adler, D.S., Kollontzi, S., Goedken, M.J., Fabris, L., Yurkow, E.J., Stapleton, P.A., 2020. Nanopolystyrene translocation and fetal deposition after acute lung exposure during late-stage pregnancy. Part. Fibre Toxicol. 17, 55. 10.1186/s12989-020-00385-9

Garcia, M.A., Liu, R., Nihart, A., El Hayek, E., Castillo, E., Barrozo, E.R., Suter, M.A., Bleske, B., Scott, J., Forsythe, K., Gonzalez-Estrella, J., Aagaard, K.M., Campen, M.J., 2024. Quantitation and identification of microplastics accumulation in human placental specimens using pyrolysis gas chromatography mass spectrometry. Toxicol. Sci. kfae021. 10.1093/toxsci/kfae021

Gatti, A.M., 2004. Biocompatibility of micro- and nano-particles in the colon. Part II. Biomaterials 25, 385–392. 10.1016/s0142-9612(03)00537-4

Gatti, A.M., Rivasi, F., 2002. Biocompatibility of micro- and nanoparticles. Part I: in liver and kidney. Biomaterials 23, 2381–2387. 10.1016/s0142-9612(01)00374-x

Gaylarde, C., Baptista-Neto, J.A., Da Fonseca, E.M., 2021. Plastic microfibre pollution: how important is clothes’ laundering? Heliyon 7, e07105. 10.1016/j.heliyon.2021.e07105

Geyer, R., Jambeck, J.R., Law, K.L., 2017. Production, use, and fate of all plastics ever made. Sci. Adv. 3, e1700782. 10.1126/sciadv.1700782

Grafmueller, S., Manser, P., Diener, L., Diener, P.-A., Maeder-Althaus, X., Maurizi, L., Jochum, W., Krug, H.F., Buerki-Thurnherr, T., Mandach, U. von, Wick, P., 2015. Bidirectional Transfer Study of Polystyrene Nanoparticles across the Placental Barrier in an ex Vivo Human Placental Perfusion Model. Environ. Health Perspect. 123, 1280–1286. 10.1289/ehp.1409271

Griffin, S., Masood, M.I., Nasim, M.J., Sarfraz, M., Ebokaiwe, A.P., Schäfer, K.-H., Keck, C.M., Jacob, C., 2017. Natural Nanoparticles: A Particular Matter Inspired by Nature. Antioxid. Basel Switz. 7, 3. 10.3390/antiox7010003

Guan, Q., Jiang, J., Huang, Y., Wang, Q., Liu, Z., Ma, X., Yang, X., Li, Y., Wang, S., Cui, W., Tang, J., Wan, H., Xu, Q., Tu, Y., Wu, D., Xia, Y., 2023. The landscape of micron-scale particles including microplastics in human enclosed body fluids. J. Hazard. Mater. 442, 130138. 10.1016/j.jhazmat.2022.130138

Hartmann, N.B., Hüffer, T., Thompson, R.C., Hassellöv, M., Verschoor, A., Daugaard, A.E., Rist, S., Karlsson, T., Brennholt, N., Cole, M., Herrling, M.P., Hess, M.C., Ivleva, N.P., Lusher, A.L., Wagner, M., 2019. Are We Speaking the Same Language? Recommendations for a Definition and Categorization Framework for Plastic Debris. Environ. Sci. Technol. 53, 1039–1047. 10.1021/acs.est.8b05297

Horvatits, T., Tamminga, M., Liu, B., Sebode, M., Carambia, A., Fischer, L., Püschel, K., Huber, S., Fischer, E.K., 2022. Microplastics detected in cirrhotic liver tissue. EBioMedicine 82, 104147. 10.1016/j.ebiom.2022.104147

Hu, J., Qin, X., Zhang, J., Zhu, Y., Zeng, W., Lin, Y., Liu, X., 2021. Polystyrene microplastics disturb maternal-fetal immune balance and cause reproductive toxicity in pregnant mice. Reprod. Toxicol. 106, 42–50. 10.1016/j.reprotox.2021.10.002

Huang, S., Huang, X., Bi, R., Guo, Q., Yu, X., Zeng, Q., Huang, Z., Liu, T., Wu, H., Chen, Y., Xu, J., Wu, Y., Guo, P., 2022. Detection and Analysis of Microplastics in Human Sputum. Environ. Sci. Technol. 56, 2476–2486. 10.1021/acs.est.1c03859

Ibrahim, Y.S., Tuan Anuar, S., Azmi, A.A., Wan Mohd Khalik, W.M.A., Lehata, S., Hamzah, S.R., Ismail, D., Ma, Z.F., Dzulkarnaen, A., Zakaria, Z., Mustaffa, N., Tuan Sharif, S.E., Lee, Y.Y., 2021. Detection of microplastics in human colectomy specimens. JGH Open Open Access J. Gastroenterol. Hepatol. 5, 116–121. 10.1002/jgh3.12457

Jambeck, J.R., Geyer, R., Wilcox, C., Siegler, T.R., Perryman, M., Andrady, A., Narayan, R., Law, K.L., 2015. Plastic waste inputs from land into the ocean. Science 347, 768–771. 10.1126/science.1260352

Jenner, L.C., Rotchell, J.M., Bennett, R.T., Cowen, M., Tentzeris, V., Sadofsky, L.R., 2022. Detection of microplastics in human lung tissue using μFTIR spectroscopy. Sci. Total Environ. 831, 154907. 10.1016/j.scitotenv.2022.154907

Jummaat, F., Yahya, E.B., Khalil H P S, A., Adnan, A.S., Alqadhi, A.M., Abdullah, C.K., A K, A.S., Olaiya, N.G., Abdat, M., 2021. The Role of Biopolymer-Based Materials in Obstetrics and Gynecology Applications: A Review. Polymers 13, 633. 10.3390/polym13040633

Kadac-Czapska, K., Knez, E., Gierszewska, M., Olewnik-Kruszkowska, E., Grembecka, M., 2023. Microplastics Derived from Food Packaging Waste—Their Origin and Health Risks. Materials 16, 674. 10.3390/ma16020674

Kosuth, M., Mason, S.A., Wattenberg, E.V., 2018. Anthropogenic contamination of tap water, beer, and sea salt. PloS One 13, e0194970. 10.1371/journal.pone.0194970

Kutralam-Muniasamy, G., Shruti, V.C., Pérez-Guevara, F., Roy, P.D., 2023. Microplastic diagnostics in humans: “The 3Ps” Progress, problems, and prospects. Sci. Total Environ. 856, 159164. 10.1016/j.scitotenv.2022.159164

Leslie, H.A., Velzen, M.J.M. van, Brandsma, S.H., Vethaak, A.D., Garcia-Vallejo, J.J., Lamoree, M.H., 2022. Discovery and quantification of plastic particle pollution in human blood. Environ. Int. 163, 107199. 10.1016/j.envint.2022.107199

Liu, S., Lin, G., Liu, X., Yang, R., Wang, H., Sun, Y., Chen, B., Dong, R., 2022a. Detection of various microplastics in placentas, meconium, infant feces, breastmilk and infant formula: A pilot prospective study. Sci. Total Environ. 854, 158699. 10.1016/j.scitotenv.2022.158699

Liu, S., Liu, X., Guo, J., Yang, R., Wang, H., Sun, Y., Chen, B., Dong, R., 2022b. The association between microplastics and microbiota in placentas and meconium: The first evidence in humans. Environ. Sci. Technol.

Locci, E., Pilia, I., Piras, R., Pili, S., Marcias, G., Cocco, P., De Giorgio, F., Bernabei, M., Brusadin, V., Allegrucci, L., Bandiera, A., d’Aloja, E., Sabbioni, E., Campagna, M., 2019. Particle Background Levels In Human Tissues—PABALIHT project. Part I: a nanometallomic study of metal-based micro- and nanoparticles in liver and kidney in an Italian population group. J. Nanoparticle Res. 21, 45. 10.1007/s11051-019-4480-y

Maher, B.A., González-Maciel, A., Reynoso-Robles, R., Torres-Jardón, R., Calderón-Garcidueñas, L., 2020. Iron-rich air pollution nanoparticles: An unrecognised environmental risk factor for myocardial mitochondrial dysfunction and cardiac oxidative stress. Environ. Res. 188, 109816. 10.1016/j.envres.2020.109816

Massardo, S., Verzola, D., Alberti, S., Caboni, C., Santostefano, M., Eugenio Verrina, E., Angeletti, A., Lugani, F., Ghiggeri, G.M., Bruschi, M., Candiano, G., Rumeo, N., Gentile, M., Cravedi, P., La Maestra, S., Zaza, G., Stallone, G., Esposito, P., Viazzi, F., Mancianti, N., La Porta, E., Artini, C., 2024. MicroRaman spectroscopy detects the presence of microplastics in human urine and kidney tissue. Environ. Int. 184, 108444. 10.1016/j.envint.2024.108444

Montano, L., Giorgini, E., Notarstefano, V., Notari, T., Ricciardi, M., Piscopo, M., Motta, O., 2023. Raman Microspectroscopy evidence of microplastics in human semen. Sci. Total Environ. 901, 165922. 10.1016/j.scitotenv.2023.165922

Munno, K., De Frond, H., O’Donnell, B., Rochman, C.M., 2020. Increasing the Accessibility for Characterizing Microplastics: Introducing New Application-Based and Spectral Libraries of Plastic Particles (SLoPP and SLoPP-E). Anal. Chem. 92, 2443–2451. 10.1021/acs.analchem.9b03626

Nie, J.-H., Shen, Y., Roshdy, M., Cheng, X., Wang, G., Yang, X., 2021. Polystyrene nanoplastics exposure caused defective neural tube morphogenesis through caveolae-mediated endocytosis and faulty apoptosis. Nanotoxicology 15, 885–904. 10.1080/17435390.2021.1930228

Ozawa, Y., Mizushima, Y., Koyama, I., Akimoto, M., Yamagata, Y., Hayashi, H., Murayama, H., 1986. Intestinal absorption enhancement of coenzyme Q10 with a lipid microsphere. Arzneimittelforschung. 36, 689–690.

Pauly, J.L., Stegmeier, S.J., Allaart, H.A., Cheney, R.T., Zhang, P.J., Mayer, A.G., Streck, R.J., 1998. Inhaled cellulosic and plastic fibers found in human lung tissue. Cancer Epidemiol. Biomark. Prev. Publ. Am. Assoc. Cancer Res. Cosponsored Am. Soc. Prev. Oncol. 7, 419–428.

Pironti, C., Notarstefano, V., Ricciardi, M., Motta, O., Giorgini, E., Montano, L., 2022. First Evidence of Microplastics in Human Urine, a Preliminary Study of Intake in the Human Body. Toxics 11, 40. 10.3390/toxics11010040

Ragusa, A., Matta, M., Cristiano, L., Matassa, R., Battaglione, E., Svelato, A., De Luca, C., D’Avino, S., Gulotta, A., Rongioletti, M.C.A., Catalano, P., Santacroce, C., Notarstefano, V., Carnevali, O., Giorgini, E., Vizza, E., Familiari, G., Nottola, S.A., 2022a. Deeply in Plasticenta: Presence of Microplastics in the Intracellular Compartment of Human Placentas. Int. J. Environ. Res. Public. Health 19, 11593. 10.3390/ijerph191811593

Ragusa, A., Notarstefano, V., Svelato, A., Belloni, A., Gioacchini, G., Blondeel, C., Zucchelli, E., Luca, C.D., D’Avino, S., Gulotta, A., Carnevali, O., Giorgini, E., 2022b. Raman Microspectroscopy Detection and Characterisation of Microplastics in Human Breastmilk. Polymers 14, 2700. 10.3390/polym14132700

Ragusa, A., Svelato, A., Santacroce, C., Catalano, P., Notarstefano, V., Carnevali, O., Papa, F., Rongioletti, M.C.A., Baiocco, F., Draghi, S., D’Amore, E., Rinaldo, D., Matta, M., Giorgini, E., 2021. Plasticenta: First evidence of microplastics in human placenta. Environ. Int. 146, 106274. 10.1016/j.envint.2020.106274

Rahman, L., Mallach, G., Kulka, R., Halappanavar, S., 2021. Microplastics and nanoplastics science: collecting and characterizing airborne microplastics in fine particulate matter. Nanotoxicology 15, 1253–1278. 10.1080/17435390.2021.2018065

Ramos, L., Berenstein, G., Hughes, E.A., Zalts, A., Montserrat, J.M., 2015. Polyethylene film incorporation into the horticultural soil of small periurban production units in Argentina. Sci. Total Environ. 523, 74–81. 10.1016/j.scitotenv.2015.03.142

Schwabl, P., Köppel, S., Königshofer, P., Bucsics, T., Trauner, M., Reiberger, T., Liebmann, B., 2019.Detection of Various Microplastics in Human Stool: A Prospective Case Series. Ann. Intern. Med. 171, 453–457. 10.7326/M19-0618

Science of Plastics [WWW Document], n.d. . Sci. Hist. Inst. URL https://sciencehistory.org/education/classroom-activities/role-playing-games/case-of-plastics/science-of-plastics/ (accessed 10.16.23).

Weingrill, R.B., Lee, M.-J., Benny, P., Riel, J., Saiki, K., Garcia, J., Oliveira, L.F.A.D.M., Fonseca, E.J.D.S., Souza, S.T.D., D’Amato, F.D.O.S., Silva, U.R., Dutra, M.L., Marques, A.L.X., Borbely, A.U., Urschitz, J., 2023. Temporal trends in microplastic accumulation in placentas from pregnancies in Hawaiʻi. Environ. Int. 180, 108220. 10.1016/j.envint.2023.108220

What is Black Carbon?, n.d. . Cent. Clim. Energy Solut. URL https://www.c2es.org/document/what-is-black-carbon/ (accessed 10.16.23).

Wick, P., Malek, A., Manser, P., Meili, D., Maeder-Althaus, X., Diener, L., Diener, P.-A., Zisch, A., Krug, H.F., Mandach, U. von, 2010. Barrier capacity of human placenta for nanosized materials. Environ. Health Perspect. 118, 432–436. 10.1289/ehp.0901200

Xue, J., Xu, Z., Hu, X., Lu, Y., Zhao, Y., Zhang, H., 2024. Microplastics in maternal amniotic fluid and their associations with gestational age. Sci. Total Environ. 920, 171044. 10.1016/j.scitotenv.2024.171044

Zhang, J., Wang, L., Kannan, K., 2020. Microplastics in house dust from 12 countries and associated human exposure. Environ. Int. 134, 105314. 10.1016/j.envint.2019.105314

Zhu, L., Zhu, J., Zuo, R., Xu, Q., Qian, Y., An, L., 2023. Identification of microplastics in human placenta using laser direct infrared spectroscopy. Sci. Total Environ. 856, 159060. 10.1016/j.scitotenv.2022.159060

